# Recapitulating apicobasal tissue polarity in extracellular matrix incorporated airway organoids

**DOI:** 10.1101/2025.09.03.672699

**Authors:** Zhuowei Gong, Dhruv Bhattaram, Laura Porritt, Kian Golestan, Amir Barati Farimani, Amy L. Ryan, Daniel J. Weiss, Xi Ren

## Abstract

The airway epithelium is a dynamic barrier that interfaces with the external environment and internal matrix along its apicobasal axis. To recapitulate this tissue arrangement in an organoid format, we present the decellularized ExtraCellular Matrix-incorporated Apical-out Airway Organoid (dECM-AoAO) that integrates basolateral matrix cues through incorporation of human lung dECM microparticles, while maintaining direct apical exposure to the exterior. Compared to the ECM-free AoAO, dECM incorporation effectively diversifies lineage distribution that better recapitulates native epithelial composition. Harnessing dECM-AoAO locomotion powered by its outward-facing ciliary beating, we developed an experimental and computational pipeline for swarm analysis of organoid group motility as a functional readout of ciliary function. Lastly, dECM-AoAO withstood cryopreservation and revival with sustained viability, lineage composition, and ciliary function, enabling future scalability and broad distribution. Together, this work establishes dECM-AoAO as a more physiologically relevant model system for investigating epithelial-ECM crosstalk during airway homeostasis, pathogenesis, and injury responses.

## 1 INTRODUCTION

The pseudostratified epithelium lining the airway lumen acts as one of the crucial respiratory defense mechanisms by maintaining barrier integrity and enabling mucociliary clearance, a process mediated by the multiciliated and secretory cells that trap and remove inhaled pathogens and foreign particles [1, 2]. Beneath the differentiated epithelium are basal cells that serve as resident epithelial stem cells of the airways which, due to their self-renewal and multilineage differentiation capacity, are essential for maintaining tissue homeostasis and replenishing the epithelium upon damage [3-6]. A defining feature of the airway epithelium is its apicobasal polarity, with the lumen-facing apical surface mediating direct interactions with inhaled environmental cues, while the basal surface adheres to the underlying extracellular matrix (ECM), enriched in fibrous proteins, proteoglycans, and glycosaminoglycans [7]. In addition to providing structural support, the ECM regulates epithelial survival, differentiation, polarization, and morphogenesis via signaling through cell-surface receptors, such as integrins and syndecan [8-11]. Thus, the development of robust airway tissue models that faithfully recapitulate such tissue architectural arrangement in terms of both cellular and extracellular composition and tissue polarity is highly desired.

Hydrogel-embedded organoid culture is a well-established strategy for exposing bioengineered airway tissues to an ECM environment. Typically, this involves embedding dissociated basal cells that are airway epithelial progenitors into an ECM hydrogel, commonly Matrigel, where each basal cell proliferates into a spheroid that can be directed to undergo mucociliary differentiation [3, 12-14]. This approach results in an airway-mimetic tissue bearing an enclosed lumen with its apical epithelial surface facing inward, referred to as Apical-in Airway Organoid (AiAO). Despite the tremendous contribution of AiAOs to our understanding of airway epithelial biology and cell-ECM interactions, the internalized apical surface makes investigation of apical epithelial processes, such as pathogen exposure or mucociliary adaptation in response to external stimuli, technically challenging. While microinjection techniques have been reported for accessing the enclosed apical-in organoid lumen, the procedure is labor-intensive and therefore has limited scalability [15].

To facilitate apical access, polarity-inverted, apical-out airway organoids (AoAOs) have been developed, allowing for direct exposure of the apical epithelial surface to environmental hazards, such as SARS-CoV-2, influenza, and enteroviruses [16, 17]. The apical-out polarity also allows the motile cilia, now presented to the organoid’s exterior, to propel whole-tissue-level AoAO motility, enabling high-fidelity visualization and assessment of ciliary (dys)function [18]. Despite these desired features, current strategies for AoAO production involve either ECM-free culture or ECM withdrawal [17], limiting its utility for modeling epithelial-ECM interactions. Further, AoAOs derived from complete ECM-free culture exhibit an oversimplified epithelial lineage composition with predominantly multiciliated cells and suffer from a lack of other key airway epithelial cell types such as secretory cells and basal cells [18]. While the underlying mechanism for this lack of epithelial diversity in AoAOs remains unclear, the absence of niche-derived ECM cues to orchestrate tissue morphogenesis may be a key contributing factor.

Beyond generic ECM materials like Matrigel, there is an increasing appreciation of biomaterials that better recapitulate organotypic ECM microenvironments for tissue engineering applications. Over the past two decades, perfusion whole-organ decellularization has been widely adopted to produce biomaterials with well-preserved native ECM composition, offering a platform for revealing the influence of ECM cues on epithelial morphogenesis under pathophysiological conditions [19, 20]. However, tissue-engineered systems involving whole-organ scaffolds are low-throughput, technically complex, and difficult to standardize. As a more tractable alternative, whole-lung dECM can be solubilized through enzymatic digestion for hydrogel production. Lung dECM hydrogels have shown beneficial effects on guiding both airway and alveolar organoid morphogenesis [21, 22]. However, organoids embedded in dECM hydrogels, much like in Matrigel-embedded culture, typically adopt an apical-in polarity, limiting direct access to the apical epithelial surface. Altogether, existing tissue-engineered systems fall short of simultaneously delivering both epithelial-ECM interaction and apical-out polarity, and a tradeoff has been necessary in terms of model selection.

To address this bottleneck, here we describe the development of an airway organoid system, termed the dECM-incorporated Apical-out Airway Organoid (dECM-AoAO), where the epithelium envelopes around a biomaterial core composed of human lung derived dECM microparticles. dECM incorporation effectively allows the organoid to recapitulate native airway tissue polarity, with the epithelium interfacing with the external environment and internal matrix along its apicobasal axis, respectively. We benchmarked the dECM-AoAO for diversifying epithelial lineage differentiation, for tracking cilia-powered organoid motility, and for compatibility with cryopreservation.

## 2 MATERIALS AND METHODS

### 2.1 Human Lung Decellularization and dECM-MP Preparation

Human lungs were acquired through the University of Vermont (UVM) Autopsy Service in accordance with institutional guidelines, and all subsequent experiments on human lungs were performed under institutionally approved protocols. No IRB approval was required to utilize autopsy specimens at UVM. Decellularization of the patient lungs utilized in this study was described and performed in a previous study according to our group’s standardized decellularization protocol [23]. In brief, whole lung lobes from patients with no history of lung disease (n=3) were decellularized through sequential perfusion of detergent and enzyme solutions through airways and vasculature utilizing a peristaltic roller pump (Stockert Shiley, SOMA Technologies) at a 2 L/min rate Sequential 2 L rinses included, 0.1% Triton-X 100 (Sigma), 2% sodium deoxycholate (SDC, Sigma), 1 M sodium chloride (NaCl, Sigma), 30 µg/mL DNase (Sigma)/1.3 mM MgSO_4_/2 mM CaCl2, 0.1% peracetic acid/4% ethanol (Sigma), followed by a deionized water wash. Confirmation of efficient decellularization for the lungs utilized in this study was performed previously, including determination of <50 ng/mg residual double-stranded DNA within decellularized lungs and the absence of DNA fragments by gel electrophoresis and nuclear staining by hematoxylin and eosin (H&E) staining, as previously described [23].

Following validation of efficient lung decellularization, lung lobes were manually dissected to isolate specific anatomical regions, including airways, vasculature, and alveolar-enriched regions [21, 24]. Dissection began at the most proximal upper airways (i.e. trachea and large airways) and vasculature (which is positioned adjacent to large airways), and progressed toward the distal regions of both the airway and vasculature trees capable of isolation. Dissected samples were subsequently lyophilized and liquid nitrogen milled into a fine powder for storage at −20ºC until future use. The isolated airway tissues were then lyophilized and cryomilled into a fine powder using liquid nitrogen milling. The resulting airway-derived dECM powder was stored at −20□°C until further use.

### 2.2 Culture of Human Bronchial Epithelial Cells (HBECs)

HBECs from healthy donors were obtained from Lonza (CC-2541). Cells were maintained at 37°C with 5% CO□ in Bronchial Epithelial Cell Growth Medium (Lonza, CC-3171) supplemented with 1□μM A83-01 (Sigma-Aldrich, SML0788), 5□μM Y-27632 (Cayman Chemical, 129830-38-2), 0.2□μM DMH-1 (Tocris, 4126), and 0.5□μM CHIR99021 (REPROCELL, 04000402). Culture vessels were pre-coated with conditioned medium from 804G cells [25].

### 2.3 Construction of dECM-Incorporated Apical-Out Airway Organoid (dECM-AoAO)

To generate dECM-AoAO, HBECs (P3-P5) were dissociated using TrypLE Express (Gibco, 12604013) and resuspended in differentiation medium consisting of PneumaCult-ALI Medium (STEMCELL Technologies, 07925) supplemented with 1□μM A83-01 and 5□μM Y-27632. Defined quantities of dECM-MP, either by weight concentration or by particle count as described in the results section, were mixed with a suspension of 1,000 HBECs, and seeded into U-bottom, 96-well, cell-repellent plates (Greiner Bio-One, 655970), where the dECM and cells were allowed to self-assemble overnight. The resulting dECM-spheroid was further differentiated for 21 days in differentiation medium without Y-27632 that was refreshed every other day. Spheroid formation and maturation were monitored throughout the culture period prior to downstream analysis.

### 2.4 Vacuum Filtration and Particle Characterization

To improve dECM-MP size distribution, dECM-MP were resuspended in Dulbecco’s Phosphate-Buffered Saline (DPBS) and passed through a Millipore Steriflip Vacuum Tube Top Filter (Millipore, SCNY00040). Brightfield images of dECM-MP before and after filtration were acquired using the EVOS M7000 Imaging System (Thermo Fisher, AMF7000). Particle count and size measurement were then performed by ImageJ. Particle size distribution analysis and histogram plotting were performed in R, including log-normal fitting. Final plots were generated using GraphPad Prism.

For fluorescence labeling, NHS (N-hydroxysuccinimide)-Cy5 (Tocris Bioscience, 5436-10) was added to the dECM-MP suspension at a final concentration of 200□µM and incubated at room temperature for 1 hour, followed by washes with DPBS to remove unbound dye. To compare dECM-MP’s intrinsic green autofluorescence and Cy5 labeling, line profile analysis was performed in ImageJ using diagonally drawn lines across dual-fluorescence images of particles, and Pearson’s correlation coefficients for fluorescence colocalization were calculated using CellProfiler. For uniform detection in immunofluorescence co-staining with cell-specific antibodies, Biotin-NHS (Millipore Sigma, H1759-5MG) was used to label dECM-MP, followed by Streptavidin-488 (Thermo Fisher, S11223) detection of biotin.

### 2.5 Organoid Imaging and Morphology Quantification

Brightfield and fluorescence images were acquired using the EVOS M7000 Imaging System. Organoid circularity and cross-sectional area were quantified from brightfield images using ImageJ. For immunofluorescence line profile analysis (Figure 3I), MATLAB was used to define the organoid center and its longest axis, along which fluorescence intensity across epithelial and particle regions was quantified.

### 2.6 Histology and Immunofluorescence Staining

For histology, dECM-AoAOs were fixed in 4% paraformaldehyde at room temperature for 30 minutes, washed thoroughly in phosphate-buffered saline (PBS), and embedded in HistoGel (Thermo Fisher Scientific, HG-4000-012) according to the manufacturer’s instructions. HistoGel-embedded organoids were processed, paraffin-embedded, and sectioned at 5-μm thickness. Sections were deparaffinized, rehydrated, and subjected to antigen retrieval by incubation in citrate-based unmasking solution (Vector Laboratories, H-3300) at 95°C for 10 minutes, followed by cooling to room temperature. After antigen retrieval, sections were permeabilized with 0.1% Triton X-100 in PBS for 10 minutes and then blocked with 1% bovine serum albumin (BSA) in DPBS for 1 hour at room temperature. Sections were subsequently incubated overnight at 4°C with mouse-anti-E-cadherin antibody (1:200, Cell Signaling Technology, 14472S), diluted in 1% BSA in DPBS, followed by incubation with Alexa Fluor 647-conjugated donkey anti-mouse IgG (1:1000, Thermo Fisher, A-31571) for 45 minutes at room temperature. Nuclei were counterstained with DAPI-containing mounting medium (SouthernBiotech, 0100-20). Images were captured using a Nikon AXR confocal microscope.

### 2.7 Whole-mount Immunofluorescence Analysis

Organoids were fixed with 4% paraformaldehyde for 1 hour at 4□°C and washed with PBS containing 0.1% Tween-20. Samples were permeabilized with 1% Triton X-100 in PBS for 45 minutes, followed by blocking with 1% BSA in PBS. Organoids were then incubated overnight at 4□°C with primary antibodies, washed, and subsequently incubated with corresponding secondary antibodies for 45 minutes at room temperature. Primary antibodies used include mouse anti-acetylated α-tubulin (Ac-α-Tub) antibody (1:200, Sigma-Aldrich, T6793), mouse anti-MUC5AC antibody (1:200, Thermo Fisher, MA5-12178), rabbit anti-CCSP (Clara Cell Secretory Protein) antibody (1:200, Thermo Fisher, PA5-78215), and mouse anti-TP63 antibody (1:100, Biocare Medical, cm163a). Secondary antibodies used include Alexa Fluor 647-conjugated donkey anti-mouse IgG (1:1000, Thermo Fisher, A-31571) and Alexa Fluor 488-conjugated donkey anti-rabbit IgG (1:1000, Thermo Fisher, R37118). Nuclei were counterstained with DAPI.

Z-stack images of stained organoids were captured using a Nikon AXR confocal microscope. For quantitative image analysis, three representative z-slices near the midsection of each organoid were captured and analyzed. To quantify surface cilia coverage, organoids stained with Ac-α-Tub and DAPI were analyzed using a convex hull approach. A convex hull was generated based on DAPI-positive nuclear pixels to define the tissue boundary and identify the organoid centroid. Each z-slice was divided into 360 one-degree angular segments radiating from the centroid. Ac-α-Tub signals located outside the convex hull, indicating surface-localized cilia, were identified within each angular segment. Morphological cilia coverage was calculated as the percentage of segments containing Ac-α-Tub signals. Quantification of TP63, CCSP, and MUC5AC expression was performed using CellProfiler. TP63 expression was calculated as the number of TP63-positive nuclei divided by the total number of DAPI-positive nuclei. CCSP and MUC5AC levels were measured as mean fluorescence intensity per slice and normalized to DAPI counts.

### 2.8 Organoid Motility Analysis and Visualization

Mature dECM-AoAOs (day 21–28) were transferred to a flat-bottom 96-well plate containing culture medium supplemented with 25□mM HEPES (Gibco, 15630080) to maintain pH stability outside of the incubator. Live imaging of organoid motility was performed at 5 frames per second using a Nikon TS2 microscope equipped with a Moment CMOS camera. To track organoid displacement over time, the Video Labeler application in MATLAB was used to draw a bounding box around each organoid in the video’s first frame, after which a Kanade-Lucas-Tomasi feature-tracking algorithm continuously shifts the bounding box location to match organoid movement through the detection of organoid features in subsequent frames. The tracking data was then converted into time-series data for the organoid centroid, from which the tracking plot was visualized and the average translational velocity over the course of the video was calculated.

### 2.9 dECM-AoAO Cryopreservation and Revival

Mature dECM-AoAOs (day 21–28) were harvested, centrifuged to remove culture medium, and resuspended in CryoStor® CS10 (STEMCELL Technologies, 07930). Organoids were transferred to cryovials and placed in a Mr. Frosty freezing container (Thermo Fisher, 5100-0001) at –80°C for 24□h prior to long-term storage in liquid nitrogen. For thawing, cryovials were rapidly warmed at 37°C water bath, washed with pre-warmed medium, and cultured for at least 24□h prior to analysis. Viability was assessed using the LIVE/DEAD™ Viability/Cytotoxicity Kit (Thermo Fisher, L3224) according to the manufacturer’s protocol. Live (Calcein-AM^+^, green) and dead (EthD-1^+^, red) cells were visualized using Nikon AXR confocal microscope. For viability quantification, Z-stack images were acquired, and three representative slices were extracted from each organoid. Image segmentation and quantification of live (green) and dead (red) areas were performed using CellProfiler and presented as the percentage of total area of live and dead signals together.

### 2.10 Statistical analysis

All quantitative data are presented as mean ± standard deviation (SD). Statistical analyses were performed using GraphPad Prism. For comparisons between two groups, unpaired two-tailed t-tests were used. For comparisons involving more than two groups with one independent variable, one-way ANOVA followed by Tukey’s multiple comparisons test was used. For experiments involving two independent variables, two-way ANOVA with Tukey’s post hoc test was applied. The Shapiro-Wilk test was used to assess the data normality. The non-parametric Kruskal-Wallis test followed by Dunn’s multiple comparisons test was used instead if normality was not met. *p*-values less than 0.05 were considered statistically significant and are indicated as follows: \**p* < 0.05, \*\**p* < 0.01, \*\*\**p* < 0.001. No data points were excluded unless justified by technical reasons. One statistical outlier was removed from Fig.□6J using a built-in Grubbs’ test (α = 0.05).

## 3. RESULTS

### 3.1 Overall workflow for dECM preparation and organoid incorporation

To establish an airway organoid platform that delivers both the apical-out polarity and native-like epithelial-ECM interaction, we developed a strategy for incorporating dECM microparticles (dECM-MP) derived from decellularized airways, obtained from autopsy specimens of patients with no known lung disease, into suspension organoid culture. To do this, whole human lungs were subjected to detergent-based perfusion decellularization to remove cellular content while preserving the native ECM [24, 26]. The airway regions were then dissected out from the decellularized lungs, lyophilized, and mechanically milled into fine powder to give rise to dECM-MP (Fig. 1A-B). Vacuum filtration was then utilized as a means to optimize particle size distribution (Fig. 1C). To incorporate dECM into organoid culture, defined quantities of dECM-MP and human bronchial epithelial cells (HBECs) were combined on a U-bottom, cell-repellent surface. The hybrid constructs (Fig. 1D) can then be directed to undergo mucociliary differentiation over a 3-week period, resulting in mature airway organoids with an apical-out polarity and basolateral dECM presence, thereafter termed dECM-AoAOs.

**Fig. 1.**
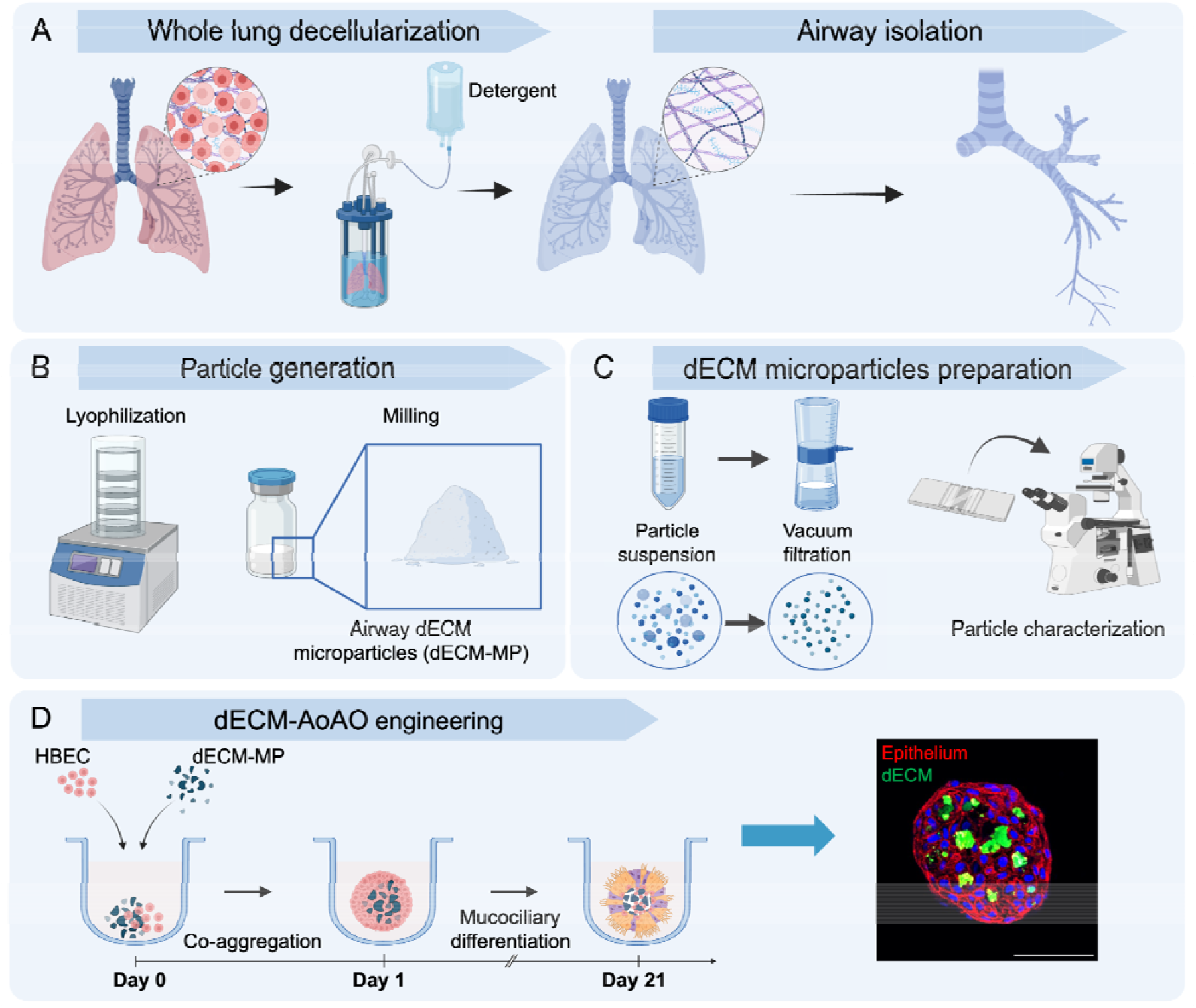
Schematic overview of dECM-AoAO engineering. (**A**) Whole-lung perfusion decellularization followed by isolation of the airway regions. (**B**) Generation of dECM-MP through decellularized tissue lyophilization and subsequently milling. (**C**) Quantification and optimization of dECM-MP particle count and size distribution. (**D**) dECM-AoAO engineering through co-aggregation of a defined quantity of dECM-MP and 1,000 HBECs, and subsequent mucociliary differentiation over three weeks of culture. Representative fluorescence image on the right showing a day-1 dECM-spheroid with constituent epithelial cells shown in red (E-Cadherin stain) and dECM-MP shown in green (visualized through the biotin labeling prior to organoid incorporation). Scale bar: 100□μm.

### 3.2 dECM-MP tracking and dose response for organoid incorporation

Fluorescence imaging performed at day 1 following the dECM-HBEC co-seeding revealed the presence of particles with green-autofluorescence within the epithelial spheroids, which was absent in stage-matched spheroids without dECM incorporation (Fig. S1), suggesting that the green-autofluorescence correlated with and indicated dECM presence. To confirm this, we performed direct fluorescence labeling of dECM-MP at a wavelength different from the green-autofluorescence via amine-reactive conjugation with the far-red Cy5 dye (Fig. 2A) and assessed the optical properties of the resulting microparticles in Dulbecco’s Phosphate-Buffered Saline (DPBS) suspension. As expected, these microparticles exhibited consistent green-autofluorescence with a high-level of colocalization with the Cy5 labeling as indicated by Pearson’s correlation analysis (Fig. 2B-D), a consistent observation across dECM-MP derived from three different donors (Fig. 2D).

**Fig. 2.**
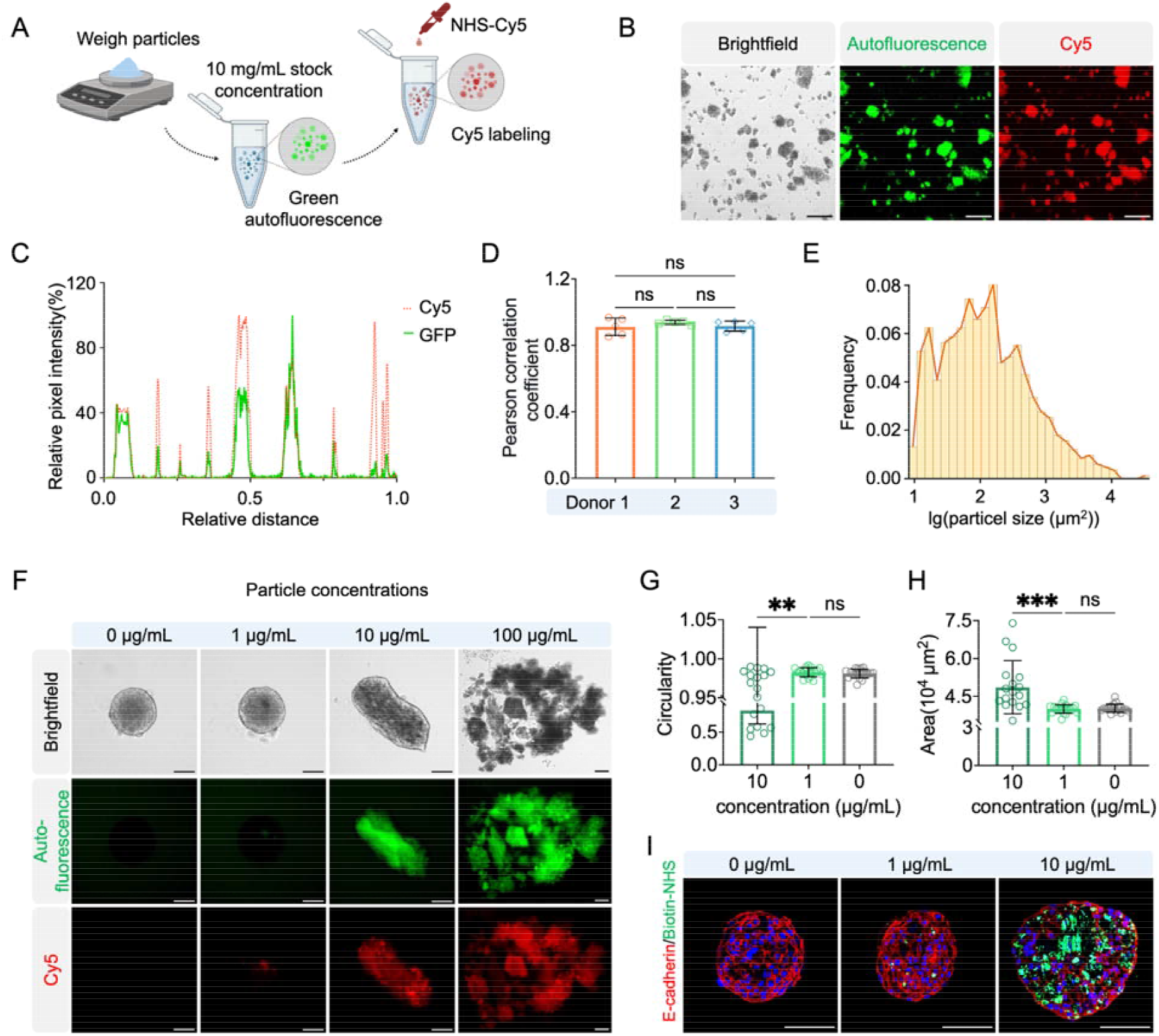
Fluorescence tracking of dECM-MP and particle dose-dependent regulation of dECM-spheroid formation. (**A**) Schematic illustrating Cy5 labeling of dECM-MP bearing intrinsic green autofluorescence. (**B**) Representative images showing brightfield and fluorescence of labeled particles. (**C**,**D**) Line profile of particle fluorescence confirms co-localization of green autofluorescence and Cy5 fluorescence (**C**), and the associated Pearson’s correlation analysis of dECM from three independent donors (**D**). Each dot represents an independent image field of dEM-MP. Data represent mean□±□SD. ns, not significant (one-way ANOVA with Tukey’s test). (**E**) Size distribution of dECM-MP displayed on a log-transformed scale. (**F**) Brightfield and fluorescence images of organoids formed with increasing dECM concentrations (0, 1, 10, and 100□μg/mL). (**G**,**H**) Quantification of organoid circularity (**G**) and cross-sectional area (**H**). Each dot represents an individual organoid. Data represent mean□±□SD. \*\*\**p*□<□0.001; \*\**p*□<□0.01; ns, not significant (Kruskal-Wallis with Dunn’s test). (I) Immunostaining of day-1 dECM-spheroids for E-cadherin (red) and biotin-labeled dECM-MP (biotin-NHS, green) at different dECM concentrations. All scale bars: 100□µm.

The mechanical milling process, while being effective in microparticle generation, resulted in dECM-MP with a broad range of size distribution (Fig. 2E), prompting us to explore whether loading based on dECM weight concentration (μg/mL) can lead to reproducible control of microparticle incorporation and consistent generation of hybrid dECM-epithelium spheroids. As control, epithelial spheroids formed in the absence of dECM (0□μg/mL) each displayed as a single compact structure with high circularity. At dECM concentration of 1□μg/mL, while the circular, compact spheroid structure was well maintained (Fig. 2G,H), tracking through the Cy5 particle labeling in live spheroid imaging revealed sparse dECM presence (Fig. 2F). An increase of dECM dose to 10□μg/mL led to much more pronounced dECM incorporation (Fig. 2F), however, the size and shape distribution of the resulting dECM-epithelium spheroids became heterogeneous (Fig. 2F-H). In line with this, further elevation of dECM dose to 100□μg/mL failed to form coherent spheroids (Fig. 2F). To further validate results observed through live organoid imaging, we turned to histological analysis and used histology-compatible biotin labeling of dECM-MP (through amine-reactive conjugation similar to Cy5 conjugation), which further confirmed robust dECM incorporation at 10□μg/mL dosage throughout the epithelium-enclosed spheroid (Fig. 2I).

### 3.3 Vacuum filtration refines dECM-MP input and improves dECM-epithelium spheroid consistency

To address the limitations encountered in our initial dECM loading efforts, we explored whether further controlling the physical properties of the input dECM-MP can improve consistency of the hybrid tissue assembly. Based on our prior dECM-MP characterization (Fig. 2B,E), we hypothesized that their broad particle size range and in particular the presence of very large particles with a size (>50 μm) on the same order as the epithelial spheroids may interfere with the consistency of dECM-epithelium assembly. To test this, we implemented a 40-µm vacuum filtration step to exclude oversized microparticles (Fig. 1C). This effectively reduced the average size and narrowed the size distribution range of the resulting filtered dECM-MP (Fig. 3A-D, Fig. S2).

**Fig. 3.**
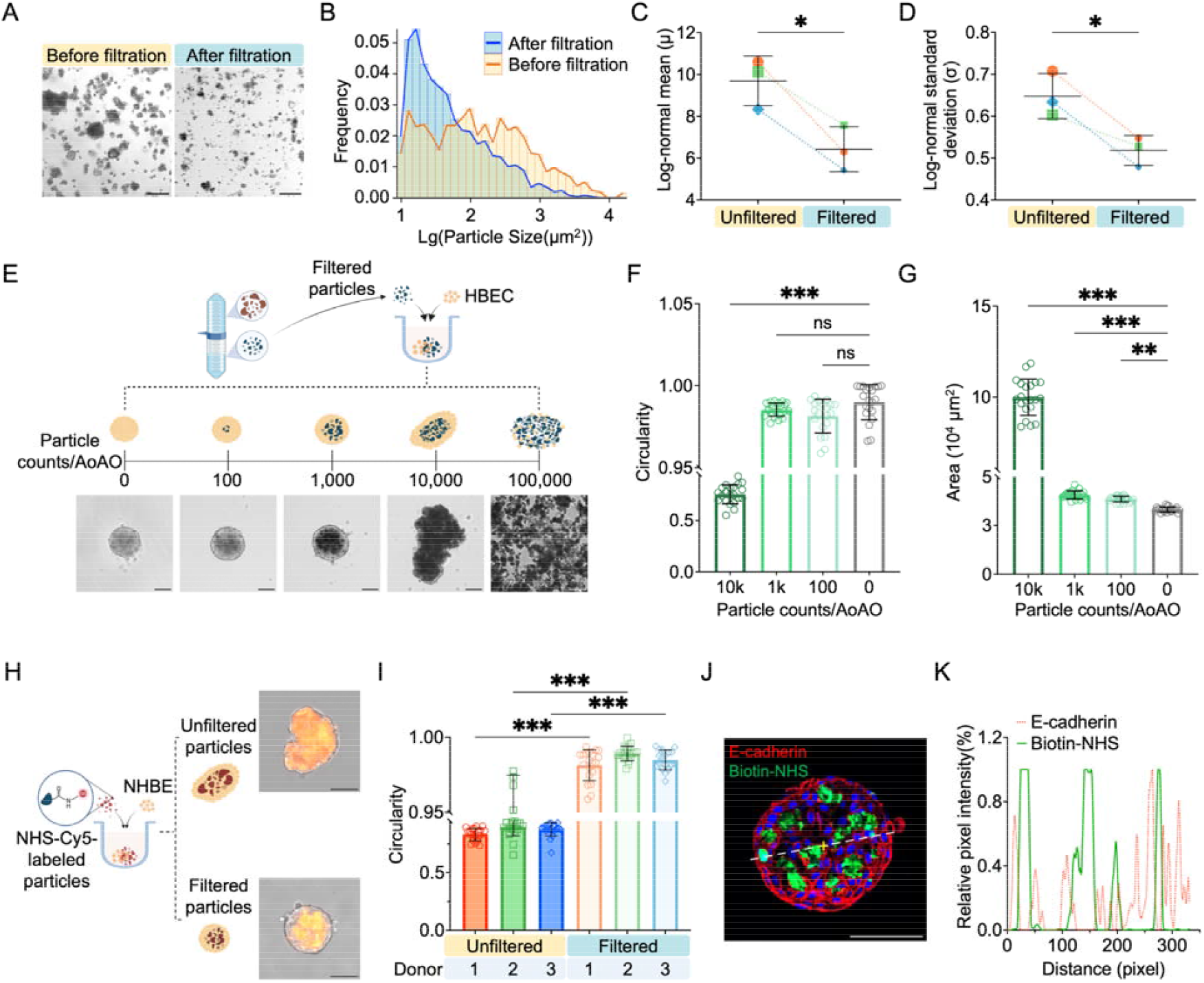
Optimization of dECM-MP size distribution and dECM-AoAO formation. (**A**) Brightfield images of dECM-MP before and after vacuum filtration. (**B**) Particle size distribution on a log-transformed scale. (**C**-**D**) Log-normal fitting of particle size confirms a significant decrease in mean (μ) and standard deviation (σ) after filtration across dECM from three donors. (**E**-**G**) Schematics and representative brightfield images (**E**) and quantification of circularity (**F**) and area (**G**) of dECM-AoAO formation using size-filtered particles, loaded at defined particle counts (0-100,000 particles per 1,000 HBECs). Each dot represents an individual organoid. (**H**-**I**) Comparison of organoids formed with filtered versus unfiltered particles (1,000 per organoid) across three donors. (**J**) Immunofluorescence staining for E-cadherin and Biotin-NHS confirms the particle incorporation in a 1,000-particle organoid (day 1). (**K**) Line profile analysis showing relative localization of E-cadherin+ cells and biotin-labeled particles. Data are shown as mean□±□SD. \*\*\**p*□<□0.001; \*\**p*□<□0.01; \**p*□<□0.05; ns, not significant (one-way ANOVA with Tukey’s test). All scale bars: 100□µm.

With improved dECM-MP size consistency, we implemented particle count-based dECM-MP loading for hybrid organoid formation. Defined numbers of filtered dECM-MP (0 to 100,000 per organoid) were co-aggregated with 1,000 HBECs to form dECM-spheroids (Fig. 3E). At high dECM-MP dosing conditions, loading with 100,000 particles per organoid completely disrupted spheroid formation (Fig. 3E); while the 10,000-particle loading still permitted tissue formation, it produced heterogeneous structures with irregular shapes (Fig. 3E-G). In contrast, hybrid spheroids formed with 100 or 1,000 particles maintained compact spherical structures with high circularity, with 1,000-particle loading producing more pronounced dECM incorporation (Fig. 3E-G, Fig. S3). Balancing the above observations, 1,000 particles per organoid was selected as the optimal condition moving forward. To further validate this strategy, we compared tissues formed from filtered versus unfiltered dECM-MP at the same particle count (1,000 per organoid) across three dECM donors, and as anticipated, dECM-epithelium spheroids derived from filtered particles consistently exhibited higher circularity (Fig. 3H,I, Fig. S4). Immunostaining of day-1 hybrid spheroids incorporating biotin-labeled dECM-MP showed multiple dECM clusters intercalated within an integrated epithelial tissue marked by E-cadherin expression (Fig. 3J,K), suggesting the epithelium acting as the main structural component holding the dECM-MP together. This full cell-matrix intermixing in the form of a cohesive and morphologically stable tissue supports our subsequent efforts to investigate cell-ECM interaction.

### 3.4 dECM incorporation diversifies airway epithelial lineage composition upon differentiation

The native airway epithelium is maintained by a coordinated balance between basal cells, secretory cells (e.g., club and goblet cells), and multiciliated cells, which together enable sustained mucociliary clearance and epithelial homeostasis. In our prior work engineering AoAOs without any ECM support, we observed an oversimplified epithelial lineage composition with most constituent cells becoming multiciliated cells and a nearly complete deprivation of any secretory cells or basal cell pool [18]. Here we probe the influence of dECM incorporation on airway epithelial lineage dynamics over the same three-week differentiation process as the AoAOs. Strikingly, in fully differentiated dECM-AoAOs, CCSP^+^ club cells and MUC5AC^+^ goblet cells can now be readily identified (Fig. 4A-C), and the TP63□ basal cells were also robustly maintained (Fig. 4A,D).

**Fig. 4.**
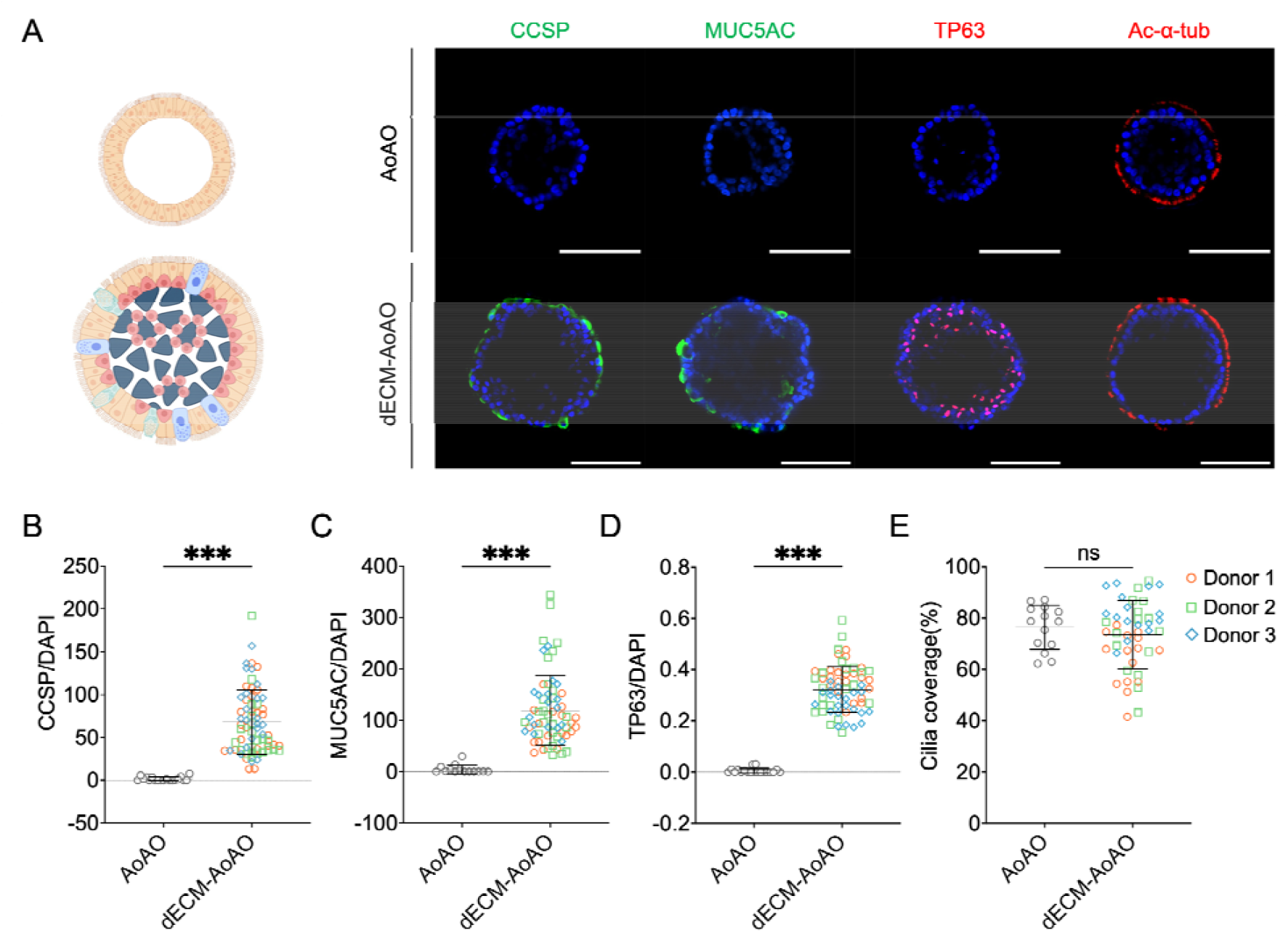
dECM incorporation enriches epithelial lineage diversity and preserves basal stem cells. (**A**) Schematic illustrations of ECM-free AoAO (top) and dECM-AoAO (bottom), along with representative immunofluorescence images showing expression of acetylated α-tubulin (Ac-α-tub; multiciliated cell), CCSP (club cell), MUC5AC (goblet cell marker), and TP63 (basal stem cell) in both organoid systems. Nuclei are stained with DAPI (blue). Scale bars: 100□μm. (**B**-**E**) Quantification of epithelial lineage marker expression in AoAOs and dECM-AoAOs: CCSP and MUC5AC expression was quantified as mean fluorescence intensity normalized to DAPI□ nuclei counts (**B**,**C**); the number of TP63□ nuclei was normalized to total DAPI□ nuclei counts (**D**); cilia coverage was calculated using morphological analysis of surface-localized Ac-α-tub signal (**E**). Each dot represents an individual organoid. Data are shown as Mean ± SD; \*\*\**p* < 0.001, ns = not significant, unpaired t-test.

Consistent with this shift, we observed a reduction in the abundance of the number of FOXJ1□ multiciliated cells in dECM-AoAOs compared to AoAOs (Fig. S5), indicating that the emergence of additional epithelial lineages occurred at the expense of the multiciliated population. To evaluate how the decrease in FOXJ1□ cell abundance may affect motile cilia presentation on organoid exterior surface, we performed confocal imaging of dECM-AoAOs stained with cilia marker Acetylated α-Tubulin (Ac-α-Tub). The outward localization of Ac-α-Tub□ cilia confirmed the apical-out epithelial polarity being well preserved following dECM-MP incorporation (Fig. 4A). We then applied a convex hull-based morphometric analysis that segmented each z-slice into 360 angular sectors from the organoid centroid and measured Ac-α-Tub signal beyond the convex boundary. Interestingly, this analysis revealed a comparable level of outer surface ciliation (73.58□±□13.33%) on dECM-AoAOs compared to that found on AoAOs (76.41□±□8.62%) (Fig. 4E), indicating epithelial diversification taking place in dECM-AoAOs without compromising the surface motile cilia coverage.

Together, our findings demonstrate dECM incorporation as an effective means to support diversified epithelial differentiation together with basal cell maintenance. This underscores the significance of enabling epithelial-ECM interaction to augment the physiological relevance of bioengineered tissue systems to deliver a closer recapitulation of not only the native lineage composition of the airway epithelium but also its interface with the external environment and internal matrix along the apicobasal axis, respectively.

### 3.5 dECM-AoAO enables functional assessment of cilia-driven tissue-level motility

We next sought to determine whether the dECM-AoAOs retained functional motile cilia, a hallmark of the healthy airway epithelium and a critical defense mechanism against inhaled pathogens and debris [27]. Here we show, when placing a fully-differentiated dECM-AoAO on a flat surface, such as that provided by a regular flat-bottom microplate, beating on its exterior-facing apical cilia could generate sufficient force to induce swirling movement of the entire organoid (Supplementary Movie S1-3). This emergent motility phenotype provides a direct, label-free readout of net, tissue-level ciliary function without the involvement of hydrogel embedding as required by our prior study [18].

To quantify motility, we tracked the displacement of individual dECM-AoAO over time and calculated their velocity as the change in position (Δx) over change in time (Δt) (Fig. 5A). This paradigm was extended to allow for simultaneous tracking of multiple dECM-AoAOs through “swarm” analysis, increasing experimental throughput without sacrificing the underlying motility phenotype (Fig. 5B). Under control conditions, mature dECM-AoAOs displayed active swirling and directional movement (Fig. 5C, top). To validate that the observed locomotion was cilia-dependent, organoids were treated with 1 mM EHNA, a dynein ATPase inhibitor that impairs ciliary beating [18]. EHNA exposure dramatically suppressed organoid motility, leading to stationary behavior and markedly shortened or absent trajectories (Fig. 5C, bottom). Quantitative analysis confirmed a significant EHNA-induced reduction in mean velocity across dECM-AoAOs produced with dECM-MP from all three donors (Fig. 5D, Supplementary Movie S4-6). Together, these results demonstrate that the dECM-AoAO platform supports robust ciliogenesis and enables real-time assessment of ciliary dynamics.

**Fig. 5.**
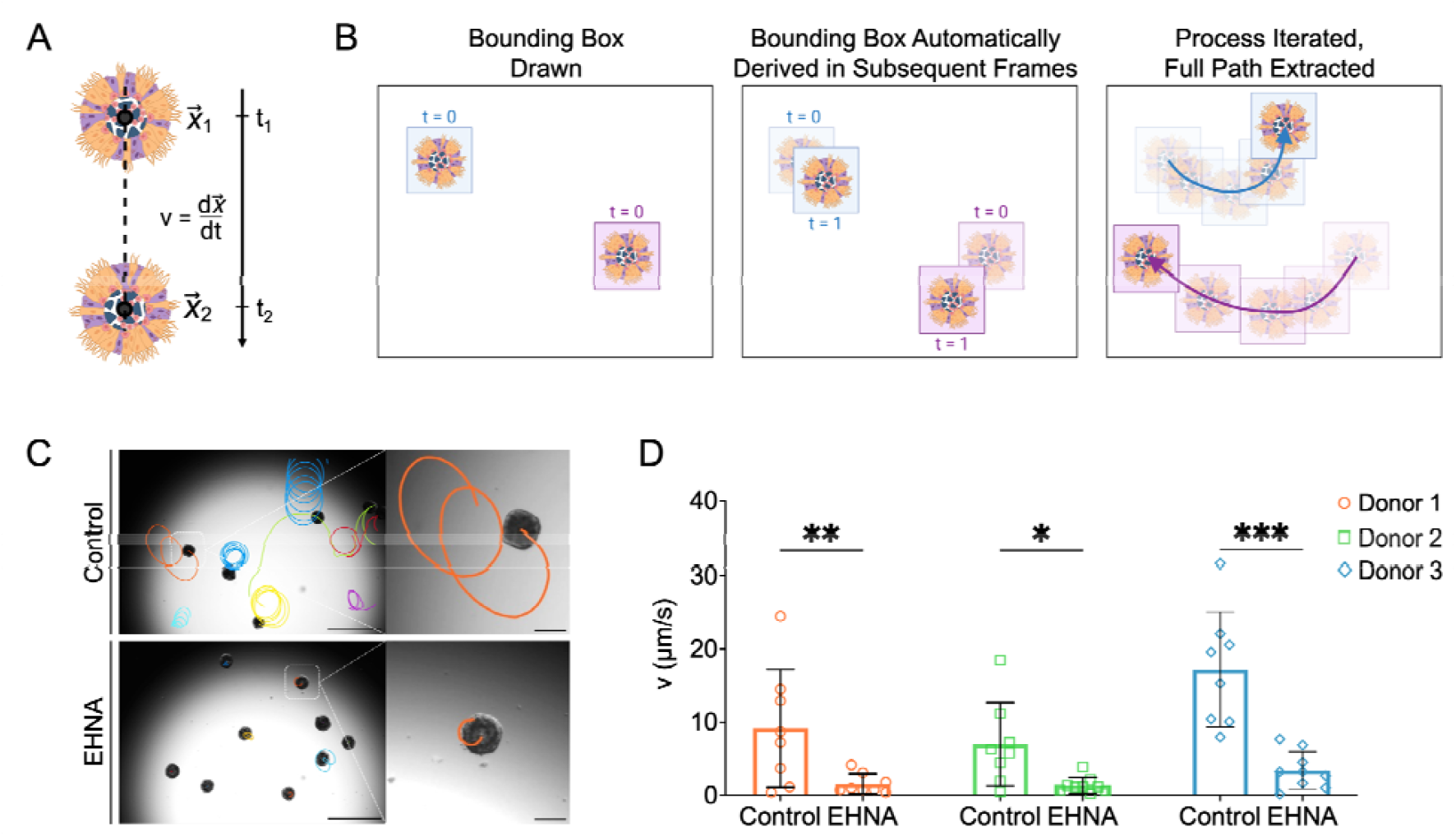
Swarm tracking dECM-AoAO 2D locomotion and its modulation by dynein inhibition (EHNA). (**A**) Schematic illustrating the computational pipeline for swarm quantification of organoid 2D locomotion. The position of each organoid is tracked at two time points (t□ and t□), and velocity (v) is calculated as the displacement over time (v = Δx / Δt). (**B**) Representative example of automated trajectory analysis. Bounding boxes are drawn around each organoid at t = 0 and then propagated frame-by-frame to extract full motility paths, allowing simultaneous tracking of multiple organoids. (**C**) Brightfield images showing representative motility patterns under control (top) and 1□mM EHNA-treated (bottom) conditions. Colored lines indicate 5-minute trajectories of individual organoids. EHNA-treated organoids showed markedly reduced movement. Scale bars (left): 500□μm; Scale bars (right): 100□μm. (**D**) Quantification of organoid velocity (µm/s) in three independent dECM donor groups. EHNA treatment significantly reduced motility compared to the control in all cases. Each dot represents an individual organoid. Bars show mean ± SD; \**p* < 0.05, \*\**p* < 0.01, \*\*\**p* < 0.001 (two-way ANOVA with Tukey’s multiple comparisons test).

### 3.6 dECM-AoAO sustained cryopreservation and revival

Much like for cells, compatibility with cryopreservation can dramatically streamline the storage, sharing, and scalable application of organoid tissue models. Such compatibility can be defined by sustained tissue viability, lineage stability, and functional properties following the freeze-thaw cycle. To assess this in the context of dECM-AoAO (Fig. 6A), we evaluated and compared epithelial viability, lineage distribution, and cilia-powered organoid motility both prior to cryopreservation and at multiple time points following revival.

**Fig. 6.**
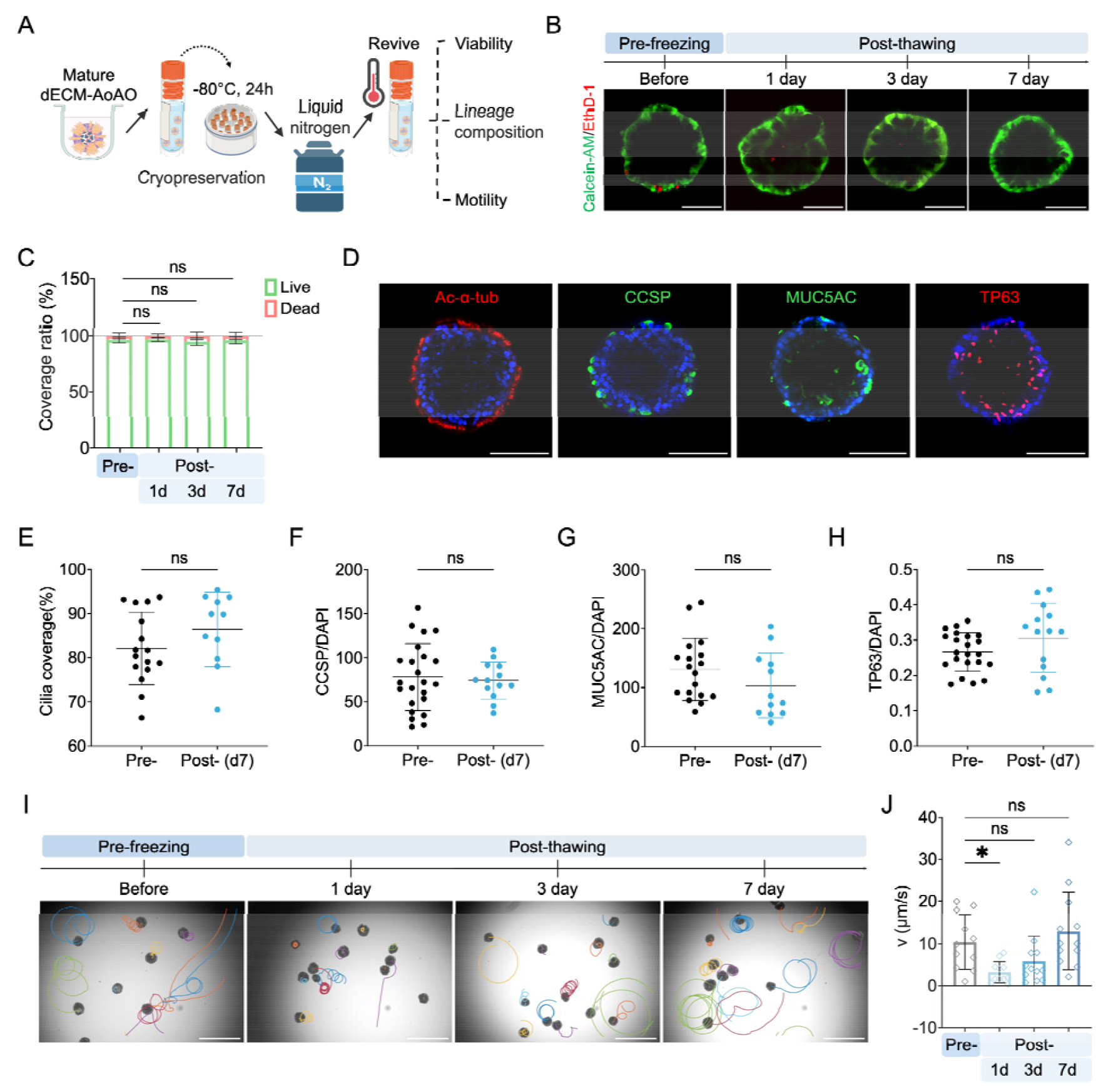
dECM-AoAO can be cryopreserved with sustained viability, lineage composition, and recoverable motility. (**A**) Schematic of cryopreservation workflow for mature dECM-AoAO. Organoids were stored in Cryostore CS10 and transferred to liquid nitrogen for cryopreservation and later revived. Data shown reflect organoids stored in liquid nitrogen for 1 day. Post-thaw assessments include cell viability, epithelial lineage composition, and cilia-powered organoid motility. (**B**,**C**) Representative Calcein-AM (live, green) and EthD-1 (dead, red) fluorescence images (**B**) and associated quantification (**C**) of organoids before freezing and at 1, 3, and 7 days after thawing. (**D**-**H**) Immunofluorescence staining (**D**) and associated quantification (**E**-**H**) of surface cilia coverage (Ac-α-tub, **E**) and the expression of CCSP (**F**), MUC5AC (**G**), and TP63 (**H**) in post-thaw organoids in comparison to pre-freeze counterparts. (**I**) Representative motility tracking images of organoids both pre-freeze and at 1, 3, and 7 days post-thaw. Colored traces indicate individual organoid trajectories. Scale bars: 500□μm. (**J**) Quantification of organoid velocity (μm/s) at each time point. Each dot represents an individual organoid. Data are presented as mean ± SD. \**p* < 0.05, ns = not significant. One statistical outlier was removed using Grubbs’ test (α = 0.05).

Live/dead staining using Calcein-AM and EthD-1 revealed that the epithelial cell viability of mature dECM-AoAO was well maintained following cryopreservation, with no significant changes in the ratio of live cell areas detected at 1, 3, or 7 days post-revival compared to that found pre-preservation (Fig. 6B,C). Further, immunofluorescence staining performed on organoids 7 days post-revival demonstrated persistent expression of key airway lineage markers, including TP63 (basal cells), CCSP (club cells), Ac-α-Tub (ciliated cells), and MUC5AC (goblet cells), with no significant alterations in their abundances compared to pre-preservation controls (Fig. 6D-H).

Upon confirming the structural integrity of dECM-AoAOs following the freeze-thaw cycle, we went on with functional assessment of cilia-powered organoid motility along the same procedure timeline. While organoid velocity was transiently reduced immediately following revival, it fully recovered to a comparable level to that observed pre-preservation within 7 days post-revival (Fig. 6I-J, Supplementary Movie S7-10). Together, these data demonstrate the full compatibility of dECM-AoAOs with cryopreservation and revival, with robust maintenance of epithelial viability, lineage diversity, and ciliary function.

## 4 DISCUSSION

In this work, we described the development of a decellularized ECM-incorporated Apical-out Airway Organoid (dECM-AoAO) system that allows seamless epithelial-ECM interaction while preserving the apical-out epithelial polarity. This model system effectively recapitulates the tissue arrangement seen in the native airway epithelium with its apical surface exposed to the external environment and its basal side interfacing with the internal matrix niche. We observed a profound influence of dECM incorporation on diversifying epithelial lineage differentiation and enabling basal cell maintenance. With surface-localized motile cilia driving tissue-level dECM-AoAO motility, we established a robust computational pipeline for swarm analysis of population locomotion on 2D surface and demonstrated its responsiveness to ciliary beating modulation. Lastly, we demonstrated dECM-AoAO’s full compatibility with cryopreservation.

The dECM-AoAO system described here combines the strengths of multiple existing bioengineered airway tissue models while addressing some of their limitations. First, dECM-AoAOs allowed more physiological incorporation of native-like extracellular environments, as it does not require the use of synthetic materials seen in air-liquid-interface culture or enzymatic solubilization of dECM for hydrogel preparation in apical-in organoid culture. In contrast to dECM hydrogel, whose properties can vary with pepsin digestion time [28], dECM-AoAOs incorporate minimally processed dECM-MP, preserving key matrix components that may be sensitive to enzymatic processing. Second, dECM-AoAOs can be engineered with high consistency through direct control of dECM-MP and cellular input, unlike the heterogeneous tissue size distribution commonly observed in apical-in organoid culture. Third, with an apical-out polarity, the dECM-AoAOs provide convenient and non-invasive access to the apical epithelial surface, facilitating future investigation of physiologic exposure to environmental hazards and apically administered therapeutics, which currently requires technically demanding interventions such as microinjection or physical disruption of the epithelial barrier [29]. Fourth, while the conventional AoAOs present an oversimplified epithelial composition [18], dECM-AoAOs demonstrate improved epithelial diversity that better mimics their native counterpart. Together, these features make dECM-AoAO a versatile and physiologically relevant model for studying human airway pathophysiology.

Our results highlight the critical role of ECM-derived cues in supporting epithelial homeostasis. Incorporating native lung dECM in the form of microparticles enables the emergence of secretory cells and preservation of the basal cell population that are otherwise depleted in matrix-free systems [18]. These findings are consistent with prior evidence that ECM components regulate airway progenitor fate via integrin-mediated signaling and matrix-bound cues [30-32]. While traditional hydrogel-embedded organoid culture can mimic certain aspects of ECM signaling, commonly used hydrogel materials, such as Matrigel derived from murine sarcoma [33, 34], are generic with little consideration of physiological relevance. Further, compared to solubilized dECM hydrogels that can indeed be prepared from specific tissues of interest, the dECM-AoAO system allows for facile incorporation of dECM in a microparticle-based format that requires minimal processing of the dECM, maximizing the preservation of pathophysiologically relevant ECM features that are fully presented to cells. By skipping the ECM solubilization procedure which can vary between enzymatic solubilization conditions, the dECM-AoAO platform delivers a higher throughput and more scalable option for recapitulating and investigating native-like ECM-epithelial interaction.

A key aspect of the AoAO model described in our previous research is an accessible assay of respiratory ciliary function through tracking the tissue-level motility generated from the exterior-facing cilia beating [18]. Here we not only demonstrate this motility assay to be compatible with the dECM-AoAO, but also have made several key technical advances. First, we removed the hydrogel embedding requirement in our prior assay format [18] by implementing a flat surface 2D locomotion assay, capturing translational velocity and, by extension, ciliary health with high sensitivity [35] and validated the correlation between organoid 2D locomotion and direct modulation of ciliary beating. The hydrogel-free organoid motility assay is significant as it enables future endeavors for whole-tissue-level ciliary functional assessment following non-invasive introduction of apical respiratory hazards and therapeutics. Furthermore, through the implementation of the “swarm” mode of the organoid locomotion analysis, we further augment the assay throughput and underscore the dECM-AoAO system’s utility for providing rapid, robust insights into ciliary health.

Another notable feature of the dECM-AoAO system is its compatibility with cryopreservation. Here we show that dECM-AoAOs can undergo a complete freeze-thaw cycle, using a standard protocol for dissociated cells, without compromising epithelial viability, lineage distribution, or ciliary function. This cryostability enhances the scalability and experimental flexibility of the tissue system by enabling batch preparation, storage, and just-in-case use. It also facilitates experimental synchronization for studies involving time-sensitive perturbations, such as infection, injury, or pharmacological treatments, and allows organoid transport across laboratories for collaborative research.

In summary, the dECM-AoAO system offers a robust and versatile tissue system that supports physiologic epithelial-ECM interaction, epithelial diversity, functional maturation, and cryopreservation. By integrating dECM-MP into a self-assembled epithelial structure, this platform captures key features of the native airway tissue architecture along its apicobasal axis. Its accessible, suspension culture-based format enables longitudinal studies, making it a valuable tool for investigating airway development, pathogenesis, and regenerative strategies. Furthermore, the simplicity of the platform allows for this model to potentially be translated into other tissues and organoid systems, including potential extensions in the intestine, liver, and pancreatic islets. Variation in cellular niches can also be coupled with variation in ECM donor, allowing for an unprecedented ability to study pathophysiological cell-ECM interactions, such as in chronic obstructive pulmonary disease, fibrosis, or cancer.

## 5 CONCLUSION

In this study, we established a decellularized lung extracellular matrix-incorporated airway organoid platform that integrates native ECM microparticles with apical-out polarity. This approach restores epithelial organization, drives the diversification of basal, ciliated, club, and goblet cell lineages, and enables label-free swarm analysis of cilia-driven motility, providing a physiologically relevant model for airway development and disease.

## Acknowledgments

We are grateful to Misti West and Garrett Struble for their support in laboratory management. We thank the Center for Biologic Imaging at the University of Pittsburgh for imaging support. We are also thankful to Dr. Hongmei Mou (Massachusetts General Hospital and Harvard Medical School) for providing the 804G cell line used in this study. All schematic figures were created with BioRender.com.

## Author contributions

ZG, DB, KG, DJW, ALR, and XR contributed to project conceptualization and methodology design. ZG, DB, LP, and KG carried out experiments and data collection, while ZG and DB performed validation, formal analysis, data curation, and visualization. ZG, DB and ABF developed the software. ZG prepared the original draft, and all authors contributed to the review and editing. ALR, DJW, and XR provided supervision, project administration, and funding acquisition.

## Data and materials availability

All data needed to evaluate the conclusions are present in the paper, Supplementary Materials, and/or the associated data repository: http://datadryad.org/share/9TUp9277z8QcuRQNhE6g_rYzAuUk-HKN3X-2YGaP27Q.

