## Supplementary material for "Recapitulating apicobasal tissue polarity in extracellular matrix incorporated airway organoids": supplementary figures.docx

Supplementary Materials

### Supplementary Figures

**Supplementary Figure S1**. Green autofluorescence emerges with dECM incorporation in airway organoids. Apical-out airway organoids (AoAOs) exhibited green autofluorescence only when co-seeded with dECM-MP. This fluorescence was absent in AoAOs without dECM, leading to the identification of the dECM particles as the source of green autofluorescence. Images were taken 1 day after seeding. Scale bar: 100 μm.

**Supplementary Figure S2.** Vacuum filtration decreases particle size and narrows the size distribution in additional donors’ dECM samples. Brightfield images and size distribution histograms of dEC-MP from Donor-2 and Donor-3 before and after filtration. Scale bars: 100 μm.

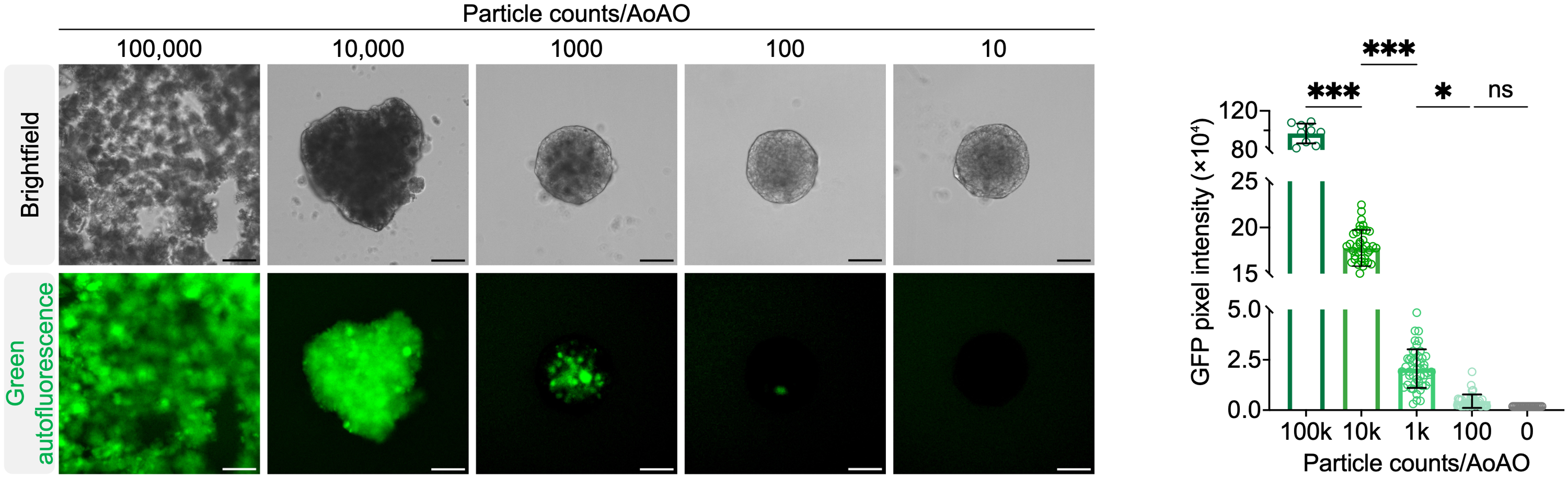

#### **Supplementary Figure S3**. GFP autofluorescence intensity confirms dose-dependent dECM incorporation in AoAOs. Brightfield and green autofluorescence images were taken 1 day after seeding. Green autofluorescence from the dECM particles was used as a readout for dECM incorporation, with higher GFP intensity observed in organoids exposed to larger particle doses. Quantification showed significantly higher fluorescence at the 1,000-particle condition compared to 100 particles, supporting greater dECM incorporation at this intermediate dose while maintaining spheroid circularity. Scale bars: 100 μm. Data are shown as mean ± SD. **p* < 0.05, ****p* < 0.001, ns = not significant (one-way ANOVA with Tukey’s test). Each dot represents an individual organoid.

**Supplementary Figure S4.** Filtered dECM particles improve spheroid morphology while retaining incorporation in additional donors. Brightfield and green autofluorescence images of AoAOs formed using filtered or unfiltered dECM-MP (1,000 particles per organoid) from Donor-2 and Donor-3. Scale bars: 100 μm.

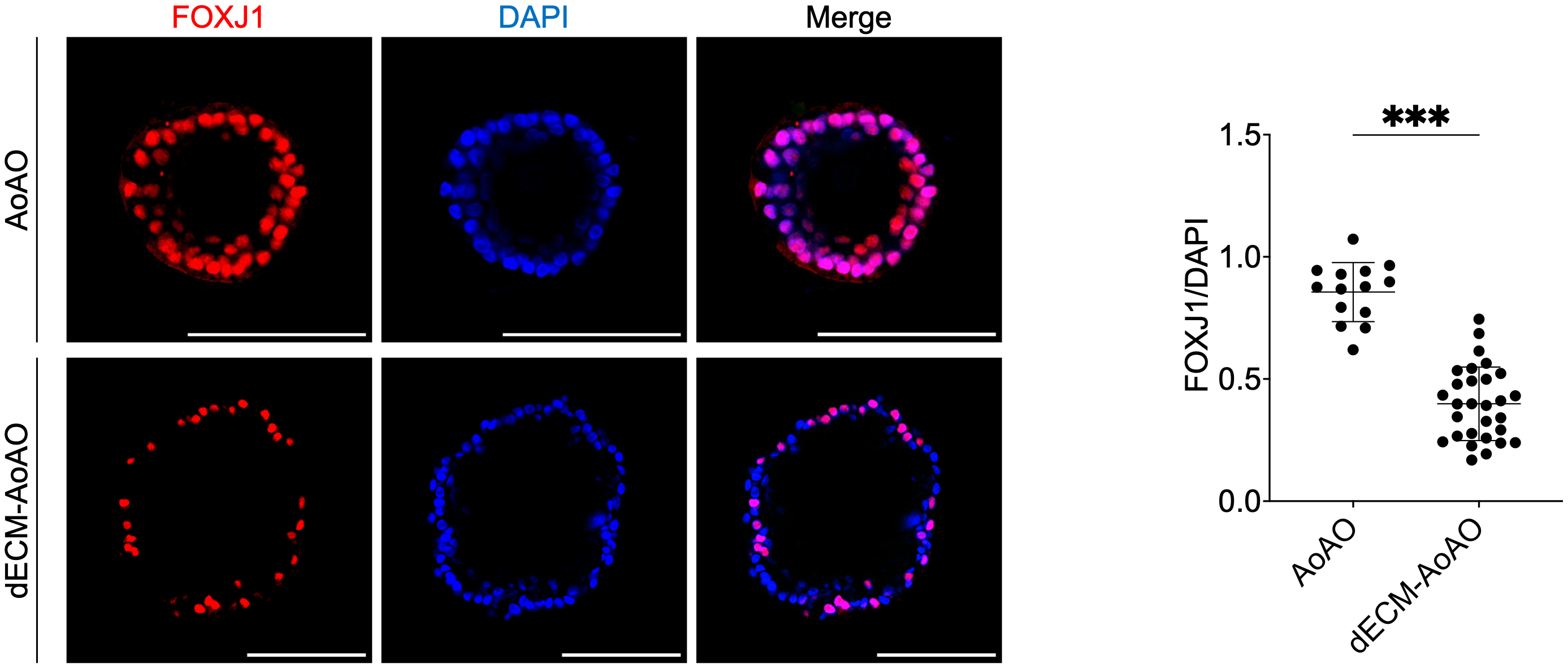

**Supplementary Figure S5.** Elevated FOXJ1 expression in AoAOs compared to dECM-AoAOs despite similar cilia coverage. Representative cross-sectional images of FOXJ1 and DAPI immunofluorescence staining in fully differentiated AoAOs and dECM-AoAOs. Quantification (right) was performed by calculating the ratio of FOXJ1⁺ cell counts to total DAPI⁺ nuclei. Each dot represents an individual organoid. Data are shown as mean ± SD; ****p* < 0.001 (unpaired t-test). Scale bars: 100 μm.

### Supplementary Movies

Supplementary Movie S1. Locomotion of donor-1 derived dECM-AOAO under control conditions. Related to Figure 5. Mature apical-out airway organoids (AOAOs) were imaged to capture their spontaneous rotational motion without EHNA treatment. Scale bar 1000 μm.

Supplementary Movie S2. Locomotion of donor-2 derived dECM-AOAO under control conditions. Related to Figure 5. Scale bar 1000 μm.

Supplementary Movie S3. Locomotion of donor-3 derived dECM-AOAO under control conditions. Related to Figure 5. Scale bar 1000 μm.

Supplementary Movie S4. Locomotion of donor-1 derived dECM AOAO after EHNA treatment. Related to Figure 5. EHNA (1 mM) was applied to mature AOAOs for 2 hours before imaging. Scale bar 1000 μm.

Supplementary Movie S5. Locomotion of donor-2 derived dECM-AOAO after 2-hour exposure to 1 mM EHNA treatment. Related to Figure 5. Scale bar 1000 μm.

Supplementary Movie S6. Locomotion of donor-3 derived dECM-AOAO after 2-hour exposure to 1 mM EHNA treatment. Related to Figure 5. Scale bar 1000 μm.

Supplementary Movie S7. Locomotion of donor-3 derived dECM-AOAO prior to cryopreservation. Related to Figure 6. Mature AOAOs were imaged before freezing to record baseline motility characteristics. Scale bar 1000 μm.

Supplementary Movie S8. Locomotion of donor-3-derived dECM-AOAO one day after thawing. Related to Figure 6. Scale bar 1000 μm.

Supplementary Movie S9. Locomotion of donor-3-derived dECM-AOAO three days after thawing. Related to Figure 6. Scale bar 1000 μm.

Supplementary Movie S10. Locomotion of donor-3-derived dECM-AOAO seven days after thawing. Related to Figure 6. Scale bar 1000 μm.
